# Germline and somatic genetic effects on gene expression and outcome in patients with Multiple Myeloma

**DOI:** 10.1101/2024.10.09.617438

**Authors:** Heini M Natri, Linh Bui, Lance Peter, Bianca Argente, Arnold Federico, Austin Gutierrez, Mei-i Chung, Jonathan Keats, Nicholas E Banovich

## Abstract

Multiple Myeloma (MM) is a hematological malignancy associated with poor prognosis and relapses. MM has a strong inherited component, and while the mutational and transcriptomic landscapes of MM have been characterized extensively, the molecular consequences of germline genetic variation in MM have not been previously explored. Here, we have leveraged matched genotype and expression data from 607 patients to map the germline and somatic regulatory variation in MM. We find that non-coding variation shapes the transcriptional landscape of MM, contributing to overall survival. These regulatory variants alter the binding of transcription factors with known importance in MM, such as IRF4, providing insights into the underlying mechanisms. These data contribute to the increasing understanding of the genetic, molecular, and cellular forces shaping MM risk and outcomes.

## Introduction

Multiple Myeloma (MM) is a hematological malignancy affecting the plasma cells in the bone marrow.^1^ In 2024, the number of new MM cases in the USA was estimated to be 35,780, accounting for 1.8% of all new cancer cases.^2^

MM predominantly affects elderly people, with a median age at diagnosis of 65 years.^3^ MM is associated with a poor prognosis, with a 5-year overall survival of 50.7% (2008-2014). While the introduction of new therapies has improved the survival rate, most patients still experience relapse.^4^ Relatives of MM patients exhibit a two-to-four fold increase in MM risk, indicating a strong inherited genetic component.^5^ MM exhibits a disparity in occurrence and mortality between the sexes and ethnicities, men and African Americans being at a higher risk than women or those of European ancestry.^6^

Genome-Wide Association Studies (GWAS) and meta-analyses have identified 23 loci associated with MM susceptibility.^7^ Many GWAS loci are located in non-coding regions and are likely to contribute to disease risk by altering gene expression.^8^ Indeed, MM risk loci are enriched in regions with active chromatin and B-cell transcription factor (TF) binding sites.^7^ In particular, variants near the transcription factor Interferon Regulatory Factor 4 (*IRF4*) are known to increase the risk of MM and other B-cell malignancies. *IRF4* is a critical regulator of the immune system, essential for plasma cell differentiation, and often overexpressed in MM.^9^ However, while the mutational and expression subtypes of MM have been extensively characterized in large-scale studies^10^, the regulatory mechanisms connecting most MM risk loci to MM occurrence, progression, and outcome remain poorly understood.

Integrated analysis of multidimensional data may improve the interpretability of human genetic variation by providing insights into specific molecular consequences of variants driving the disease process. In particular, expression quantitative trait loci (eQTL) analyses can be used to link GWAS risk loci to phenotypes by identifying the target genes of regulatory variants. Such molecular drivers represent excellent candidates for mechanistic disease biomarkers and can be targeted therapeutically in individuals at high genetic risk. Here, we have applied eQTL analyses to discover germline genetic variants and non-coding somatic mutations that modulate gene expression and disease outcome in a large cohort of patients with MM.

## Materials and methods

### CoMMpass data collection and processing

Peripheral blood whole genome sequence and tumor whole transcriptome sequence data were available for 607 donors who participated in the Multiple Myeloma Research Foundation’s (MMRF) CoMMpass study (release IA15).^11^ Tumor whole genome sequence data was available for 519 donors. Samples were collected and processed as described in Skerget et al. 2024.^10^

### Read mapping and gene expression level quantification

Whole genome and RNA short sequence reads were aligned to a custom reference genome containing the human reference hs37d5, human ribosomal complete repeating unit sequence, genome sequences of 14 oncoviruses, and FASTA sequences of ERCC RNA spike-in mixes.

WGS reads were aligned using *BWA-MEM* v.0.7.8^12^ BAM files were processed with *Picard*’s *MarkDuplicates* (http://broadinstitute.github.io/picard) and *GATK IndelRealigner*^*13*^ with default settings.

RNAseq reads were aligned using *STAR* v.2.3.1z_r395^14^ with the following parameter settings: outFilterMismatchNmax 10, outFilterMismatchNoverLmax 0.1, alignIntronMin 20, alignIntronMax 1000000, alignMatesGapMax 1000000, alignSJoverhangMin 8, alignSJDBoverhangMin 1, seedSearchStartLmax 30, chimSegmentMin 15, chimJunctionOverhangMin 15. Read counts were quantified using *HTseq*.^15^

### Multi-dimensional scaling and differential expression analysis of male and female tumors

We used multi-dimensional scaling and differential expression analyses to detect transcriptome-wide differences between male and female tumors. Gene expression data was filtered to retain genes with a mean FPKM >0.5 in at least one of the sexes. For differential expression, the filtered, untransformed count data were normalized and logCPM transformed with *voom*.^16^ A design matrix with sex as a predictor variable and biobank, batch, and age as covariates was used to fit the linear model. DEGs between comparisons were identified with limma^17^ by computing empirical Bayes statistics with the eBayes function. An FDR-adjusted p-value threshold of 0.01 and an absolute log2 fold-change threshold of 2 were used to select significant DEGs. For multi-dimensional scaling, transformed expression data was adjusted to account for covariates using the removeBatchEffect function of limma. Multi-dimensional scaling was done using the plotMDS function of limma. Euclidean distances between pairs of samples were calculated based on 100 genes with the largest standard deviations between samples.

### Variant calling, imputation, and filtering

Germline variants were called by first producing per-sample raw genotype-likelihoods using *GATK HaplotypeCaller* with a minimum base quality score 10, and then joint genotyping all the per-sample gVCFs using *GenotypeGVCFs*.^13,18^

Imputation of missing genotypes was performed in 5Mb intervals across chromosomes with *IMPUTE2*^*19*^, using the 1000 Genomes Phase 3 multiethnic reference panel. Imputation of the X-chromosomal alleles was carried out by treating the pseudoautosomal regions (PAR1 and PAR2) as autosomal and adding the *--chrx* flag for the imputation for the X-linked region. Genotype data were filtered to retain genotypes with probability scores ≥0.9 and variants with minor allele frequency (MAF) >0.05 and minor allele count (MAC) >10.

Somatic mutations were called from paired tumor and normal samples using Mutect^20^ as described in Skerget et al. 2024.^10^

### Gene expression data normalization and accounting for non-genetic sources of variation

Gene expression data were filtered and normalized for eQTL analyses. Genes with FPKM >0.1 and a read count of >6 in at least 50 samples were considered expressed and included in the analysis. The distributions of FPKM in each sample and gene were transformed to the quantiles of the standard normal distribution.

In addition to germline and somatic genetic effects, copy number alterations (CNAs) may alter gene expression levels and cause spurious signals in eQTL mapping. We used linear regression to model the expression levels of each gene as a function of the copy number state of each individual in the said gene. The residuals of these models were used as the final expression data. CNA data was available for 574 individuals and 25,553 of the 25,602 tested genes. The copy-number state of the remaining individuals/genes was set to 0.

100 PEER factors were included as covariates to account for technical batch effects in the expression data (Supplementary Materials). Sex, as well as five genotype Principal Components (PCs) were included as covariates to account for population structure.

### Germline *cis*-eQTL mapping

Germline variant effects on tumor gene expression were identified by linear regression as implemented in *QTLtools*.^21^ Variants within 1Mb of the gene under investigation were considered for testing. *p*-values of top-associations adjusted for the number of variants tested in *cis* were obtained using 10,000 permutations. False discovery rate (FDR) adjusted *p*-values were calculated to adjust for multiple phenotypes tested. Significant associations were selected using an FDR adjusted *p*-value threshold of 0.01.

### Identifying sex-specific effects of germline *cis*-eQTLs

As the effects of genetic variants on gene expression levels may differ between the sexes^22,23^, we used stratified analyses to detect sex-specific eQTLs. To identify high-confidence instances of differential eQTL effects between the sexes, we identified *cis*-acting germline eQTLs in males and females separately, as well as in the joint analysis of both sexes. eQTLs that were called in one sex but not in the other or in the joint analysis were called as sex-specific.

### Somatic *cis*-eQTL mapping

Single nucleotide mutations in 520 tumor samples were annotated using SnpEff.^24^ To identify non-deleterious, non-coding, putatively regulatory mutations, the data was filtered to exclude mutations that alter the gene product by retaining variants with a SnpEff effect impact classification “MODIFIER.” For somatic eQTL mapping, clusters of mutations within 50bp were binned to identify recurrently mutated loci. Loci with mutations in fewer than 5 tumor samples were removed from further analyses. The center position of each bin was used in the eQTL analysis. Associations between mutations within a 50kb *cis*-window of TSSs and target gene expression levels were obtained with *QTLtools* using the nominal pass. Significant associations were selected based on an FDR-adjusted *p*-value threshold 0.2.

### Genomic context of *cis*-eQTLs

To understand the genomic context of the putative regulatory variants detected in the eQTL analyses, eQTL loci were annotated using the R package *annotatr*. ^25^

### Transcription factor binding site motif analysis

To test whether the regulatory variants detected in the eQTL analyses disrupt the binding of known transcription factors (TFs), we used *HOMER*^*26*^ to analyze eQTL positions for enriched TF binding site motifs. Sets of background positions (germline) and regions (somatic) were generated as follows: First, the distribution of the distances between significant germline eQTLs and the transcription start sites (TSSs) of the target genes was obtained. Then, a number of random genes corresponding to the number of significant eQTLs in the unstratified analysis were obtained, and sites matching the distance distribution were used as background sites. *findMotifsGenome*.*pl* with the default region size 200bp was used for *de novo* motif discovery and to detect enriched known motifs among the germline eQTLs.

For the somatic analysis, a set of background regions was created similarly to the germline analysis, using the somatic eQTL bin sizes. *findMotifsGenome*.*pl* was run with the given bin sizes. *HOMER* uses a hypergeometric test to test for the enrichment of known binding motifs derived from published ChIP-seq experiments that are optimized for motif degeneracy thresholds.

### Co-localization of known MM GWAS risk loci and eQTL loci

Locations of a total 161 unique previously published GWAS loci, including 49 associated with MM ^7,27–29^, 25 associated with MM with IgH translocations^30^, 14 with MM with hyperdiploidy^30^, 9 with MM and monoclonal gammopathy^28^, 1 with MM survival^31^, 20 with B-cell malignancies including MM^32^, 54 with clostridium difficile infection in multiple myeloma^33^, and 4 with Bortezomib-induced peripheral neuropathy in MM^34^ were obtained from the EBI GWAS catalog. Without access to genome-wide summary statistics, we were unable to perform formal colocalization analyses. Alternatively, we aimed to identify eQTL loci that are in close linkage (R^2^>0.8) with the published GWAS loci. We used the LDlink^35^ LDproxy tool to identify tightly linked loci surrounding the GWAS loci. R^2^ values were obtained based on the 1000 genomes European populations (CEU, TSI, IBS, GBR, FIN).

### Survival analysis

We used the Cox’s proportional hazards model as implemented in the *coxph* function of the R package *survival* (https://github.com/therneau/survival) to test for the effects of eGene expression on overall and progress-free survival in the CoMMpass cohort. We used a penalized spline method as implemented in the function *pspline* to estimate nonlinear relationships in the model. FDR-adjusted *p*-values were calculated with the *p*.*adjust* function from the *stats* package in base R.

### Cell lines and cell culture

The MM cell lines used in this study are male cell lines (OCI-My5 and LP-1) and female cell lines (KMS20 and AMO1). Cells were cultured at 37°C with 5% CO_2_ in advanced RPMI 1640 (Gibco) supplemented + Glutamax supplemented with 4% Fetal Bovine Serum and 1% penicillin-streptomycin. HEK293T cells used for lentiviral production were cultured at 37°C with 5% CO_2_ in DMEM supplemented with 4 mM L-Glutamine, 10% Fetal Bovine Serum and 1% penicillin-streptomycin.

### Open chromatin profiling by ATAC-seq

Sex-specific effects of eQTLs may be due to sex differences in the patterns of chromatin accessibility. We used open chromatin profiling of male and female-derived MM cell lines to identify regions of differential chromatin accessibility between the sexes and to more generally characterize the chromatin landscape in MM. For ATAC-seq, transposition and library construction were performed as described in.^36^ 100,000 cells from cell cultures with high viability (above 90%) were treated with DNAse (Worthington Cat# LS002007) at a concentration of 200U/ml in culture medium at 37°C for 30 minutes. Cells were washed thoroughly 3 times with 1X PBS to remove DNAse completely, and the cell pellet was collected by centrifuging at 500 RCF at 4°C for 5 minutes. Cell lysis was performed for 3 minutes on ice using 50 *μ*l of cold ATAC-resuspension buffer (RSB) containing 0.1% NP40, 0.1% Tween-20 and 0.01% Digitonin. Lysis buffer was washed out with 1ml of cold ATAC-RSB containing 0.1% Tween-20 but no NP40 or digitonin, and the nuclei pellet was collected by centrifuging at 500 RCF at 4°C for 10 minutes. Transposition reaction was performed using 25 *μ*l 2X TD buffer, 2.5 *μ*l transposase (Illumina) for 30 minutes at 37°C in a thermocycler with 1000 RPM mixing. Transposed fragments were cleaned up with Zymo DNA Clean and Concentrator-5 kit (Zymo Research, Cat# D4014) and pre-amplification of transposed libraries was performed with 2X NEBNext Master Mix (NEB, Cat#) using the following program: 72°C for 5 min, 98°C for 30 sec, and 5 cycles of (98°C for 10 sec, 63°C for 30 sec, 72°C for 1 min) with the corresponding primers (Supplementary Table 3). 5 *μ*l of pre-amplification products were used in a qPCR reaction to determine the additional cycles needed and transposition libraries were purified using Zymo

DNA Clean and Concentrator-5 kit (Zymo Research, Cat# D4014). Library QC was assayed on Agilent 2200 TapeStation using D5000 high-sensitivity tape, and library quantification was performed on Qubit prior to sequencing on NextSeq High 500/550 platform (Illumina).

### ATAC-seq processing and differential chromatin accessibility analysis

ATAC-seq reads were mapped to the custom human reference genome using bowtie2 v. 2.3.0.^37^ SAM files were converted to BAM with samtools^38^ and coordinate-sorted with the Picard toolkit’s *SortSam*.^39^ BAM files of the same samples from different lanes were merged and mitochondrial reads were removed using samtools. Optical duplicates were removed with Picard toolkit’s *MarkDuplicates*. Fragment distribution statistics were collected with Picard toolkit’s *CollectInsertSizeMetrics*. Peak calling was performed using HMMRATAC^40^ with default parameters. Peak annotation and plots were generated using the R package ChIPseeker.^41^ Differentially accessible regions between male and female cell lines were called using edgeR^42^.

## Results

### Germline genetic effects on tumor gene expression MM

In the joint analysis of both sexes, we detected 5,983 *cis*-eQTLs with an FDR-adjusted *p*-value <0.01 (Fig. 1A). Furthermore, we detected 3,394 and 1,484 eQTLs in the male and female-specific analyses, respectively (Fig. 1A, B). In total, we detected 9,282 unique eQTLs associated with 6,137 eGenes. We detected a total of 8,801 eSNPs, most of which were located in non-coding regions (Fig. 1C). Effect sizes ranged from -1.72 to 1.45 with an average slope of 0.32 (Fig. 1E). The average distance from TSS was 64.5 kb (Fig. 1F); most cis-eQTLs were located near transcription start sites (TSSs), with 34.6% of all eQTLs across the joint and sex-specific analyses being located within 20kb of TSSs. In the ATACseq analysis of 3 male and 3 female-derived MM cell lines, we detected 36,241 regions with an accessible chromatin structure; 9.5% of the eQTLs detected here were located within these accessible regions.

**Figure 1.**
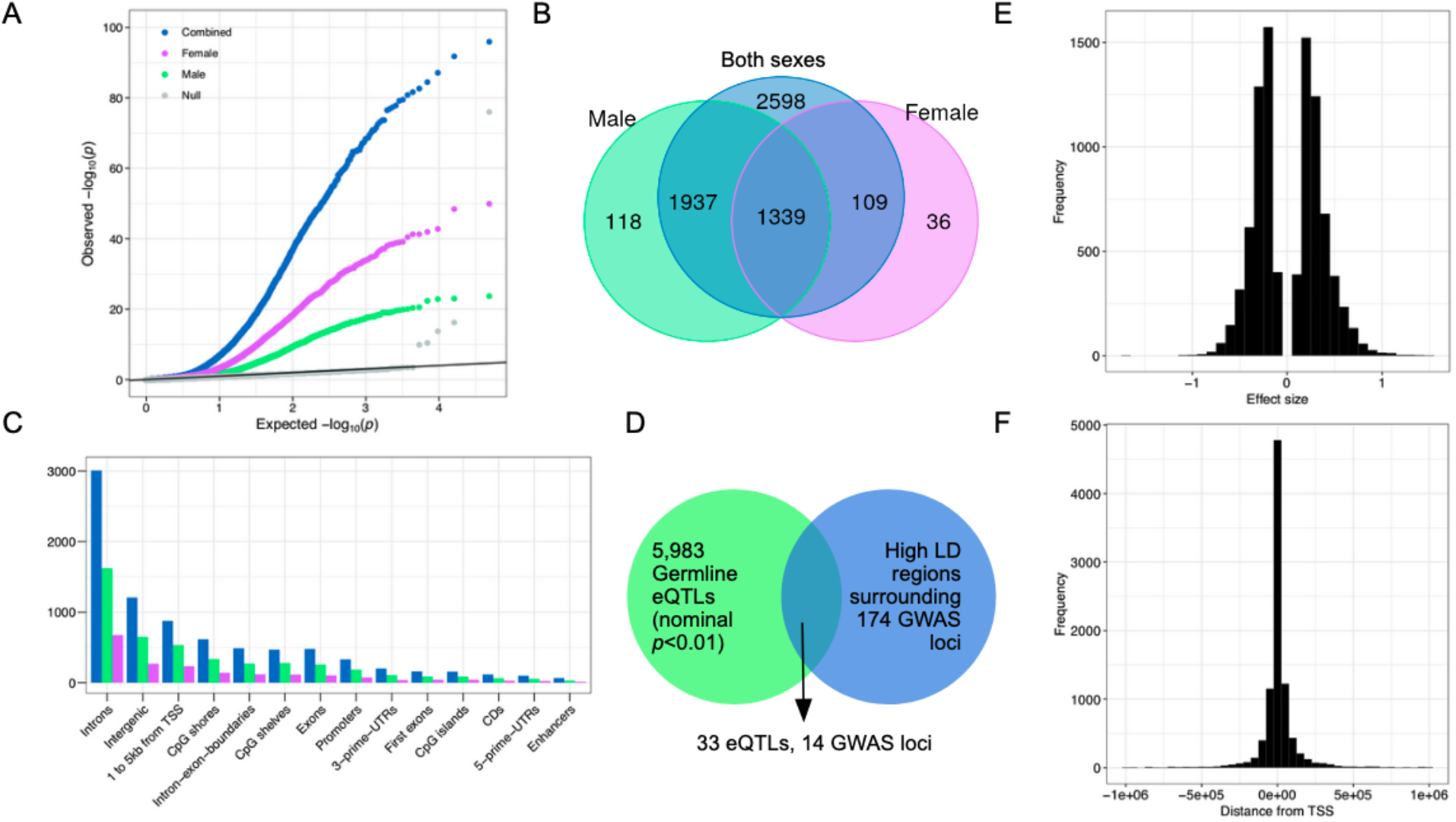
**A:** A QQ-plot of the combined and sex-stratified germline eQTL calls as well as a null-distribution based on shuffled data. **B**: Overlap of eGenes detected in combined and sex-stratified analyses. **C**: Genomic annotations of germline eQTLs detected in the combined and sex-stratified analyses. **D**: Overlap of germline eQTLs and high LD regions surrounding MM risk GWAS loci. **E-F**: Histograms of germline eQTL effect sizes and germline eSNP distances from TSS.

**Figure 2.**
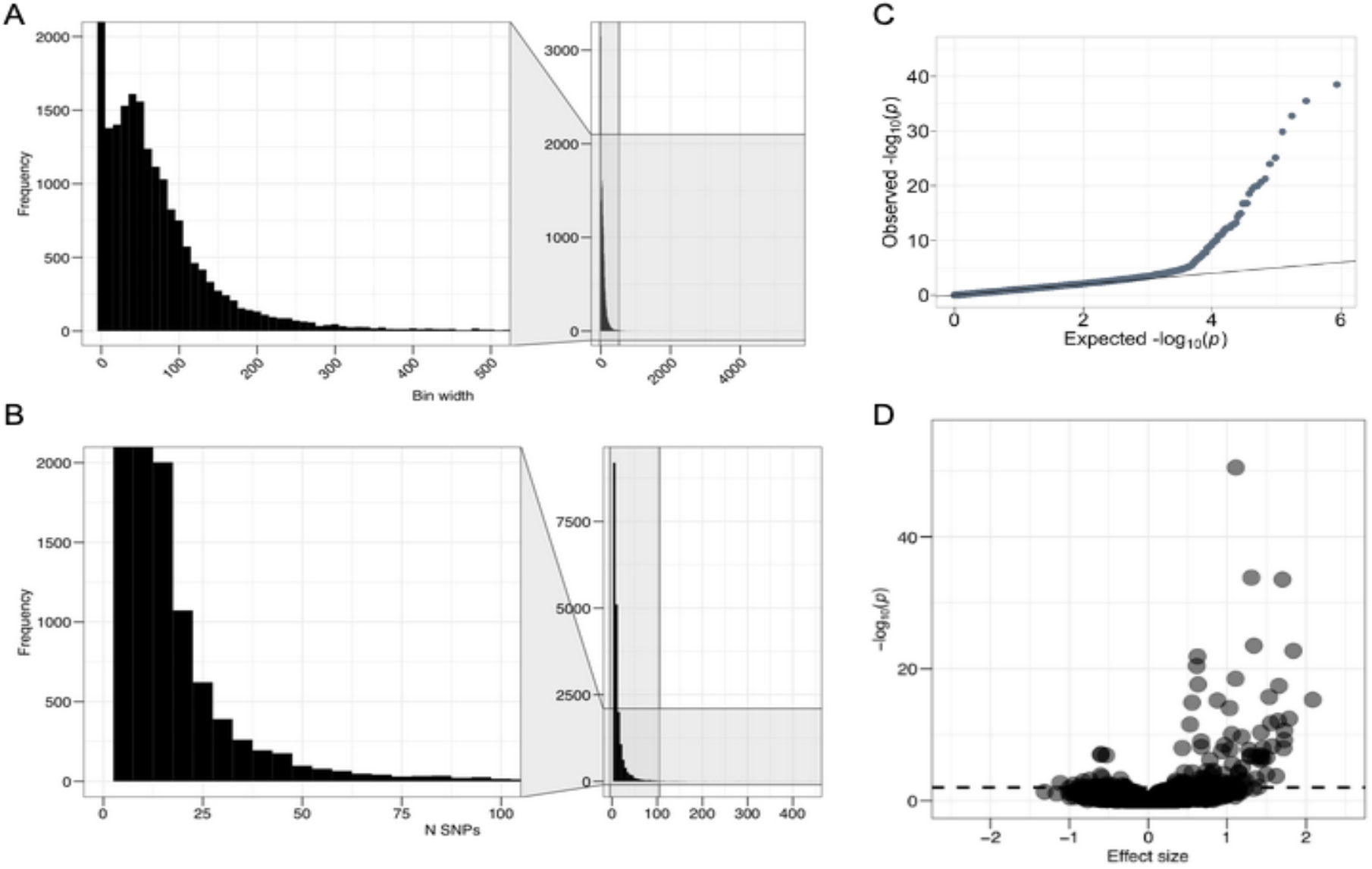
**A, B:** Histograms of the somatic bin widths and numbers of mutations in bins. **C:** A QQ-plot of the somatic eQTLs. **D:** A volcano plot of the somatic eQTL effect sizes and -log10 *p*-values.

To connect the regulatory loci detected here to MM risk, we examined the overlap of germline eQTL and high-LD regions surrounding 174 MM related GWAS loci (Fig. 1D). We find 33 eQTL in high LD with GWAS risk variants associated with MM and MM related clinical phenotypes, such as clostridium difficile infection.

### Sex-specific effects of regulatory variants

We used sex-stratified analyses and an interaction model to detect sex-specific regulatory effects of germline variants. Out of the total 8,801 unique variants detected in the joint and sex-stratified analyses, 11.85% exhibited sex-specific effects. 2,190 eQTLs were detected only in the male analysis and not in the female analysis nor in the joint analysis of both sexes. Similarly, 1,089 eQTLs were detected only in the female analysis. We further inspected genes that were under *cis*-regulatory control in only one of the sexes but not in the other or in the joint analysis of both sexes and detected 118 male-specific and 36 female-specific eGenes. Concordant with previous studies^22,23^, these genes were not differentially expressed between the sexes, indicating that the observed sex-specific regulatory effect is not due to overall expression differences between the sexes, but likely due to differences in chromatin accessibility and/or transcription factor activity.

In the ATACseq analysis, we detected 5,716 regions that exhibited differential chromatin accessibility between the male and female-derived cell lines (FDR-adjusted *p*<0.1). However, due to limited ATACseq data, limited power, and the overall low number of sex-specific eQTLs, we find no enrichment of sex-specific regulatory variants in these regions.

### Non-coding somatic mutations shape tumor gene expression in MM

Non-coding somatic mutations were binned to define recurrently mutated loci (Methods). On average, the 19,631 loci used in somatic eQTL mapping contained 12.95 mutations (min 5, max 439) and spanned 72.94bp (min 1bp, max 5,579bp). With a q-value threshold of 0.2, we detected 266 somatic eQTLs (mutation and gene pairs) associated with 188 recurrent mutations and 208 target genes. Somatic mutation calls on variable sites of the genome might indicate false positives, however, only three of the somatic eQTLs overlapped germline variants in this cohort. This network of somatic eQTLs was disrupted in 499 (96%) out of the 519 patients, indicating that non-coding regulatory somatic mutations have a wide impact in MM. Of the loci used in the somatic eQTL analysis, 4,638 loci (23.6%) contain NCVs reported in the COSMIC database. In total, 15,159 COSMIC NCVs are located within these 4,638 loci. When examining the 266 somatic eQTLs called with a *q*-value threshold of 0.2, 74 loci (28%) contained NCVs (in total 1,678 NCVs) in COSMIC.

### Effects of regulatory variants and mutations on transcription factor binding

We used a TF binding motif enrichment analysis to investigate whether the detected eQTLs are enriched in transcription factor binding sites. We find that germline eQTLs alter the binding of 8 known TF binding motifs (Benjamini *q*-value <0.05). Furthermore, using a *de novo* motif discovery method, we find 37 additional enriched TF binding motifs, including *IRF4*.

### Regulatory variants and mutations impact overall survival in the CoMMpass cohort

To connect regulatory variants to outcomes, we employed Cox’s hazards ratio to test for the effects that the expression levels of these genes have on overall and progress-free survival. 681 (%) of all eGenes had a significant (adj. *p*<0.01) effect on overall survival. These include genes associated with canonical regulatory pathways with significant in MM, such as *IRF4* (Fig. 4A, B).

**Figure 4.**
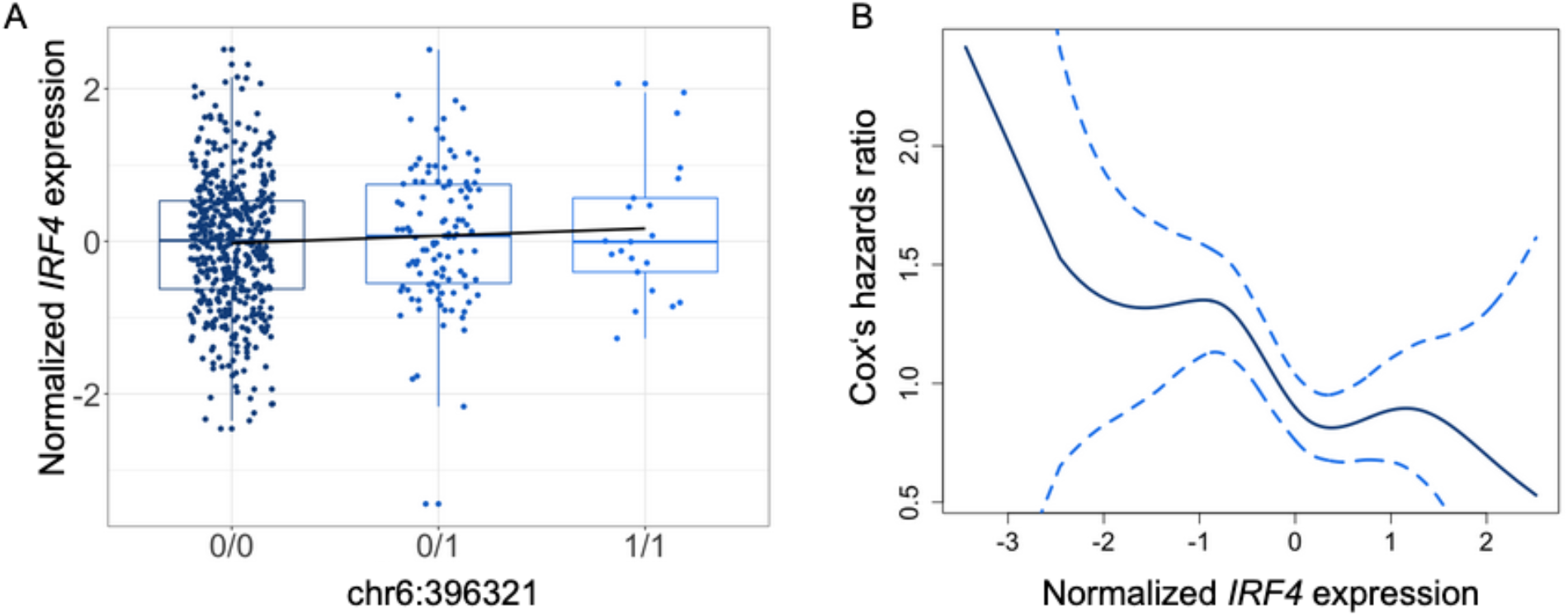
**A**, Normalized expression of IRF4 across genotypes in chr6:396321. **B**, Cox’s hazards ratio with 95% CI for the effect of normalized IRF4 expression on overall survival.

## Conclusions

Here, we have identified germline variants and non-coding somatic mutations that regulate the transcriptional landscape of MM. We identify regulatory variants that alter the expression and binding of transcription factors of biological significance, such as *IRF4*, a master regulator of an aberrant gene expression in MM.^9^ Gaining insight into the mechanisms underlying these regulatory loci, we identify a subset of eQTL residing on open chromatin regions in MM cell lines. Connecting the expression levels of the target genes of germline and somatic eQTLs to outcomes, we identify a number of regulatory variants associated with overall survival in MM. Together, these findings contribute to the genetic, molecular, and transcriptomic subtyping and characterization of MM and provide a useful resource for future studies.

## Supporting information

TFBS enrichment results, somatic

TFBS enrichment results, germline

## Data availability

Genome-wide eQTL summary statistics and ATACseq data are available on FigShare: https://doi.org/10.6084/m9.figshare.26307886.v1

## Acknowledgments

We wish to that the Multiple Myeloma Research Foundation and the participants of the CoMMpass study, who made this study possible.

