## Supplementary material for "Germline and somatic genetic effects on gene expression and outcome in patients with Multiple Myeloma": TFBS enrichment results, somatic

/scratch/hnatri/CoMMpass/TFBS\_motif\_analysis/HOMER\_genome\_sites\_with\_bg\_somatic\_q020/ - Homer de novo Motif Results


### Homer *de novo* Motif Results (/scratch/hnatri/CoMMpass/TFBS\_motif\_analysis/HOMER\_genome\_sites\_with\_bg\_somatic\_q020/)

Known Motif Enrichment Results  
Gene Ontology Enrichment Results  
If Homer is having trouble matching a motif to a known motif, try copy/pasting the matrix file into
STAMP  
More information on motif finding results: HOMER
| Description of Results
| Tips
  
Total target sequences = 179  
Total background sequences = 78  
\* - possible false positive  

|  |  |  |  |  |  |  |  |  |
| --- | --- | --- | --- | --- | --- | --- | --- | --- |
| Rank | Motif | P-value | log P-pvalue | % of Targets | % of Background | STD(Bg STD) | Best Match/Details | Motif File |
| 1 | G T A C C G A T A C T G C T A G C T A G C A T G T A C G G C T A T C A G G C A T | 1e-39 | -8.984e+01 | 23.46% | 2.32% | 53.2bp (50.8bp) | WT1(Zf)/Kidney-WT1-ChIP-Seq(GSE90016)/Homer(0.768) More Information \| Similar Motifs Found | motif file (matrix) |
| 2 | G C A T A T C G T A G C G C T A A T G C T A C G G A C T A T C G T G C A T A G C | 1e-36 | -8.356e+01 | 22.35% | 0.81% | 58.6bp (8.7bp) | Arntl/MA0603.1/Jaspar(0.712) More Information \| Similar Motifs Found | motif file (matrix) |
| 3 | A T G C G T A C T C G A A T C G G A T C G A T C A C G T A T G C G T A C C T G A A T C G G T A C | 1e-36 | -8.356e+01 | 22.35% | 0.97% | 56.6bp (0.0bp) | ZNF415(Zf)/HEK293-ZNF415.GFP-ChIP-Seq(GSE58341)/Homer(0.587) More Information \| Similar Motifs Found | motif file (matrix) |
| 4 | A G T C G T A C T G C A C A T G A T C G C G A T A T G C C G T A T A C G A T C G A T C G T A G C | 1e-34 | -8.047e+01 | 21.79% | 2.32% | 56.3bp (38.3bp) | Nr5a2/MA0505.1/Jaspar(0.635) More Information \| Similar Motifs Found | motif file (matrix) |
| 5 | G C A T C G A T A C G T C G A T A G T C C G T A A T C G C G A T C A T G C T G A G T C A G T C A | 1e-33 | -7.741e+01 | 21.23% | 2.52% | 58.4bp (18.3bp) | Stat6/MA0520.1/Jaspar(0.688) More Information \| Similar Motifs Found | motif file (matrix) |
| 6 | T G C A G T C A T A G C G T A C G C A T C T A G T C G A T G C A | 1e-32 | -7.518e+01 | 26.26% | 3.02% | 59.9bp (1.4bp) | SIX1/MA1118.1/Jaspar(0.798) More Information \| Similar Motifs Found | motif file (matrix) |
| 7 | T C A G T C G A T A G C C G T A G A T C C A T G C A G T C A G T T A C G T G A C | 1e-32 | -7.438e+01 | 20.67% | 1.85% | 53.1bp (34.4bp) | Creb3l2/MA0608.1/Jaspar(0.629) More Information \| Similar Motifs Found | motif file (matrix) |
| 8 | A T C G C G T A A T G C G A T C G C A T A T C G C A G T T A C G C A G T A G C T | 1e-28 | -6.496e+01 | 24.02% | 3.50% | 54.7bp (20.5bp) | POL009.1\_DCE\_S\_II/Jaspar(0.752) More Information \| Similar Motifs Found | motif file (matrix) |
| 9 | C T A G C T A G A C T G T A G C G A T C C G A T A C T G C T A G C T G A A G T C G A T C A T G C | 1e-25 | -5.976e+01 | 17.88% | 1.74% | 59.6bp (17.0bp) | POL011.1\_XCPE1/Jaspar(0.632) More Information \| Similar Motifs Found | motif file (matrix) |
| 10 | C T G A C G A T A C T G G A C T C A G T A C T G T G A C G C A T C T A G A C G T G C T A C G A T | 1e-25 | -5.783e+01 | 26.26% | 3.95% | 56.9bp (35.3bp) | PB0122.1\_Foxk1\_2/Jaspar(0.646) More Information \| Similar Motifs Found | motif file (matrix) |
| 11 | A C T G C T A G C A G T C A T G C A T G C T A G C G A T A G C T A G T C G A T C | 1e-24 | -5.695e+01 | 17.32% | 0.99% | 48.5bp (0.0bp) | ZNF692(Zf)/HEK293-ZNF692.GFP-ChIP-Seq(GSE58341)/Homer(0.651) More Information \| Similar Motifs Found | motif file (matrix) |
| 12 | A C G T A T G C A G T C A G T C C A T G T C A G A T C G T C A G A G T C G A T C | 1e-24 | -5.695e+01 | 17.32% | 1.26% | 58.1bp (0.0bp) | Zfx/MA0146.2/Jaspar(0.643) More Information \| Similar Motifs Found | motif file (matrix) |
| 13 | C G A T A C T G C G T A A C G T A C T G C T G A G T A C C G T A | 1e-24 | -5.565e+01 | 25.70% | 4.11% | 52.2bp (37.2bp) | Tbx20(T-box)/Heart-Tbx20-ChIP-Seq(GSE29636)/Homer(0.816) More Information \| Similar Motifs Found | motif file (matrix) |
| 14 | A T G C A G T C G A C T C A T G A T C G C A G T A T C G A C G T A G T C G A T C T G A C C G A T | 1e-23 | -5.417e+01 | 16.76% | 0.44% | 57.3bp (0.0bp) | ZNF165(Zf)/WHIM12-ZNF165-ChIP-Seq(GSE65937)/Homer(0.604) More Information \| Similar Motifs Found | motif file (matrix) |
| 15 | G C A T T A C G G C A T A T C G A T G C G A T C G T A C C A G T | 1e-22 | -5.286e+01 | 21.23% | 2.76% | 56.5bp (3.6bp) | THAP1/MA0597.1/Jaspar(0.751) More Information \| Similar Motifs Found | motif file (matrix) |
| 16 | A G C T C G A T A G T C G T C A A C G T C T G A G A T C C G T A C T A G C G T A | 1e-22 | -5.144e+01 | 16.20% | 1.84% | 51.3bp (46.0bp) | PB0028.1\_Hbp1\_1/Jaspar(0.687) More Information \| Similar Motifs Found | motif file (matrix) |
| 17 | A C T G C G T A C T A G C A T G T A G C G T A C G T A C C T G A A C T G C G T A A C T G C A T G | 1e-22 | -5.144e+01 | 16.20% | 1.39% | 58.1bp (0.0bp) | Nr2f6(var.2)/MA0728.1/Jaspar(0.590) More Information \| Similar Motifs Found | motif file (matrix) |
| 18 | A C G T A C G T A C G T A G T C A G T C G A T C G A C T A G T C G C T A C A G T A C G T G C A T | 1e-21 | -4.874e+01 | 15.64% | 1.59% | 54.3bp (19.6bp) | NFATC3/MA0625.1/Jaspar(0.660) More Information \| Similar Motifs Found | motif file (matrix) |
| 19 | A T G C G C A T C T A G T A C G C T G A G T A C G T C A A G C T | 1e-20 | -4.721e+01 | 23.46% | 4.17% | 56.7bp (45.2bp) | ZNF416(Zf)/HEK293-ZNF416.GFP-ChIP-Seq(GSE58341)/Homer(0.770) More Information \| Similar Motifs Found | motif file (matrix) |
| 20 | A G C T A C G T A C T G A T C G C A G T T G C A A G T C T A G C | 1e-20 | -4.721e+01 | 23.46% | 4.05% | 58.2bp (43.8bp) | PH0166.1\_Six6\_2/Jaspar(0.693) More Information \| Similar Motifs Found | motif file (matrix) |
| 21 | G C T A C G T A G C A T C T G A C G A T G C T A A G T C C G A T G A T C G A T C | 1e-19 | -4.599e+01 | 19.55% | 3.77% | 51.6bp (8.3bp) | PB0174.1\_Sox30\_2/Jaspar(0.636) More Information \| Similar Motifs Found | motif file (matrix) |
| 22 | T C G A A G T C A G T C C G T A A T C G T G C A A T C G A G T C T G C A A T G C | 1e-18 | -4.317e+01 | 22.35% | 4.30% | 59.7bp (1.0bp) | RUNX-AML(Runt)/CD4+-PolII-ChIP-Seq(Barski\_et\_al.)/Homer(0.627) More Information \| Similar Motifs Found | motif file (matrix) |
| 23 | G A T C G T A C C A T G C T A G C A T G G A T C G C A T A T C G C G T A A T C G | 1e-18 | -4.230e+01 | 25.14% | 5.31% | 62.7bp (33.9bp) | Zfp809(Zf)/ES-Zfp809-ChIP-Seq(GSE70799)/Homer(0.647) More Information \| Similar Motifs Found | motif file (matrix) |
| 24 | G A C T G T A C G A T C G T C A G A C T G T A C G A T C T C G A A G C T A G T C G A T C T C G A | 1e-17 | -4.090e+01 | 13.97% | 2.00% | 52.7bp (25.2bp) | Egr1(Zf)/K562-Egr1-ChIP-Seq(GSE32465)/Homer(0.680) More Information \| Similar Motifs Found | motif file (matrix) |
| 25 | C G A T G T A C G C T A A T G C C G T A T A C G G C T A A T C G A C G T T G A C G C T A A C G T | 1e-17 | -3.944e+01 | 17.88% | 3.30% | 56.6bp (53.9bp) | FOS/MA0476.1/Jaspar(0.593) More Information \| Similar Motifs Found | motif file (matrix) |
| 26 | C A G T A T C G A G C T A G T C G T A C A C T G C A G T C T A G A C T G A G T C | 1e-14 | -3.348e+01 | 12.29% | 2.15% | 57.7bp (17.5bp) | SF1(NR)/H295R-Nr5a1-ChIP-Seq(GSE44220)/Homer(0.723) More Information \| Similar Motifs Found | motif file (matrix) |
| 27 | G C A T A C G T A C G T A C T G A G C T A C G T C A T G A C G T C A T G T C G A A G C T C T A G | 1e-14 | -3.348e+01 | 12.29% | 2.04% | 47.0bp (52.7bp) | PB0121.1\_Foxj3\_2/Jaspar(0.720) More Information \| Similar Motifs Found | motif file (matrix) |
| 28 | A C G T C G T A A G T C A C G T A C T G A C G T A G T C A G T C | 1e-14 | -3.322e+01 | 16.20% | 3.63% | 57.7bp (41.6bp) | PB0050.1\_Osr1\_1/Jaspar(0.682) More Information \| Similar Motifs Found | motif file (matrix) |
| 29 | C T A G C A T G A C T G A C T G A C G T G A C T G A C T A C G T A C G T C A G T | 1e-13 | -3.184e+01 | 18.99% | 3.95% | 42.3bp (37.5bp) | ZNF384/MA1125.1/Jaspar(0.727) More Information \| Similar Motifs Found | motif file (matrix) |
| 30 | T C A G T G C A T A G C C G T A A T C G C G A T C A G T C A T G C A G T T C G A | 1e-13 | -3.123e+01 | 15.64% | 2.71% | 54.9bp (25.0bp) | MYB/MA0100.3/Jaspar(0.752) More Information \| Similar Motifs Found | motif file (matrix) |
| 31 | C T A G T C G A C G A T T C A G C G A T A C T G C A T G A T C G C G A T T A C G G C T A C G A T | 1e-13 | -3.123e+01 | 15.64% | 3.81% | 47.5bp (0.3bp) | ETS:RUNX(ETS,Runt)/Jurkat-RUNX1-ChIP-Seq(GSE17954)/Homer(0.630) More Information \| Similar Motifs Found | motif file (matrix) |
| 32 | T G C A A G T C G A C T A T G C G T A C G C A T A G T C G T A C T A G C G C A T A T G C G C A T | 1e-13 | -3.110e+01 | 11.73% | 0.00% | 55.8bp (0.0bp) | ZSCAN22(Zf)/HEK293-ZSCAN22.GFP-ChIP-Seq(GSE58341)/Homer(0.720) More Information \| Similar Motifs Found | motif file (matrix) |
| 33 | T G A C A G T C C G T A G T A C A G T C C G T A A G C T A G T C C G T A A G T C A G T C C G T A | 1e-12 | -2.877e+01 | 11.17% | 2.36% | 53.0bp (6.6bp) | YY2/MA0748.1/Jaspar(0.583) More Information \| Similar Motifs Found | motif file (matrix) |
| 34 | C G T A C G A T C G A T A C G T A C T G G C T A A T G C C G T A C G T A C G A T A C G T C G A T | 1e-12 | -2.877e+01 | 11.17% | 2.43% | 50.9bp (36.9bp) | Pax7(Paired,Homeobox),longest/Myoblast-Pax7-ChIP-Seq(GSE25064)/Homer(0.638) More Information \| Similar Motifs Found | motif file (matrix) |
| 35 | A C T G A C T G C G T A A G T C C G T A A G T C A C G T A G T C C G T A A C T G | 1e-12 | -2.785e+01 | 22.91% | 6.76% | 60.5bp (93.7bp) | Nkx2-5(var.2)/MA0503.1/Jaspar(0.694) More Information \| Similar Motifs Found | motif file (matrix) |
| 36 \* | G C A T C G A T C G A T G C T A C A G T C A G T G C T A A G C T C G A T G C A T G C A T G C A T | 1e-11 | -2.650e+01 | 10.61% | 1.86% | 53.1bp (24.8bp) | CDX1/MA0878.1/Jaspar(0.745) More Information \| Similar Motifs Found | motif file (matrix) |
| 37 \* | G T A C A C G T A C G T G A T C A G T C C A G T A C G T A C G T | 1e-10 | -2.494e+01 | 21.79% | 7.14% | 53.2bp (36.7bp) | NFATC3/MA0625.1/Jaspar(0.735) More Information \| Similar Motifs Found | motif file (matrix) |
| 38 \* | G A C T G A T C G T A C G A C T A G T C T G A C G A C T A G C T G A T C A G T C G A T C G A C T | 1e-10 | -2.429e+01 | 10.06% | 0.62% | 42.1bp (0.0bp) | EWSR1-FLI1/MA0149.1/Jaspar(0.612) More Information \| Similar Motifs Found | motif file (matrix) |
| 39 \* | C A G T C T A G T C A G C T G A A C T G G C T A G T A C G T C A T A G C G T C A A T G C G C A T | 1e-10 | -2.429e+01 | 10.06% | 1.97% | 55.8bp (43.4bp) | Bapx1(Homeobox)/VertebralCol-Bapx1-ChIP-Seq(GSE36672)/Homer(0.570) More Information \| Similar Motifs Found | motif file (matrix) |
| 40 \* | C T A G T A G C G C A T C G T A T G A C C G A T T G C A C T A G | 1e-9 | -2.103e+01 | 17.88% | 5.10% | 52.2bp (40.6bp) | PB0154.1\_Osr1\_2/Jaspar(0.719) More Information \| Similar Motifs Found | motif file (matrix) |
| 41 \* | A T G C A G T C C G T A A T C G A T C G A T G C C G A T A C T G C T A G A C G T T A C G G T C A | 1e-8 | -1.850e+01 | 11.73% | 3.03% | 58.4bp (40.7bp) | ZEB1/MA0103.3/Jaspar(0.619) More Information \| Similar Motifs Found | motif file (matrix) |
| 42 \* | A T G C A T G C G T C A A G C T A C T G T A G C G T C A A G C T | 1e-6 | -1.418e+01 | 7.26% | 2.49% | 56.6bp (27.2bp) | Pit1(Homeobox)/GCrat-Pit1-ChIP-Seq(GSE58009)/Homer(0.714) More Information \| Similar Motifs Found | motif file (matrix) |
