## Supplementary material for "Germline and somatic genetic effects on gene expression and outcome in patients with Multiple Myeloma": TFBS enrichment results, germline

/scratch/hnatri/CoMMpass/TFBS\_motif\_analysis/HOMER\_genome\_sites\_with\_bg\_wCNVs\_joint/ - Homer de novo Motif Results


### Homer *de novo* Motif Results (/scratch/hnatri/CoMMpass/TFBS\_motif\_analysis/HOMER\_genome\_sites\_with\_bg\_wCNVs\_joint/)

Known Motif Enrichment Results  
Gene Ontology Enrichment Results  
If Homer is having trouble matching a motif to a known motif, try copy/pasting the matrix file into
STAMP  
More information on motif finding results: HOMER
| Description of Results
| Tips
  
Total target sequences = 5101  
Total background sequences = 5190  
\* - possible false positive  

|  |  |  |  |  |  |  |  |  |
| --- | --- | --- | --- | --- | --- | --- | --- | --- |
| Rank | Motif | P-value | log P-pvalue | % of Targets | % of Background | STD(Bg STD) | Best Match/Details | Motif File |
| 1 | G C A T G C T A A T C G G T C A T G C A C G T A G T A C G T A C C G T A A C T G G T A C A G T C | 1e-58 | -1.346e+02 | 0.90% | 0.03% | 54.0bp (35.9bp) | IRF4(IRF)/GM12878-IRF4-ChIP-Seq(GSE32465)/Homer(0.649) More Information \| Similar Motifs Found | motif file (matrix) |
| 2 | G A C T A T C G A C T G C G A T C G A T G C T A G C T A C T G A G C T A C G A T C G A T A C T G | 1e-45 | -1.045e+02 | 0.74% | 0.03% | 55.8bp (0.0bp) | PB0109.1\_Bbx\_2/Jaspar(0.649) More Information \| Similar Motifs Found | motif file (matrix) |
| 3 | T G C A G T C A C G T A A C G T A T C G G A C T C G T A G A T C C G T A C G A T A C T G G T C A | 1e-39 | -9.002e+01 | 0.67% | 0.03% | 56.6bp (45.5bp) | Oct6(POU,Homeobox)/NPC-Pou3f1-ChIP-Seq(GSE35496)/Homer(0.658) More Information \| Similar Motifs Found | motif file (matrix) |
| 4 | A C G T A C T G A C T G A G T C A C G T A C T G A C G T A G C T C G T A A C T G A T G C G T C A | 1e-39 | -9.002e+01 | 0.67% | 0.03% | 45.6bp (47.9bp) | NFIA/MA0670.1/Jaspar(0.588) More Information \| Similar Motifs Found | motif file (matrix) |
| 5 | G A T C C G T A A C G T C G A T G C A T A G C T G A T C C G A T A C T G G T C A C G A T A C T G | 1e-36 | -8.296e+01 | 0.63% | 0.04% | 50.4bp (18.0bp) | EWS:ERG-fusion(ETS)/CADO\_ES1-EWS:ERG-ChIP-Seq(SRA014231)/Homer(0.711) More Information \| Similar Motifs Found | motif file (matrix) |
| 6 | G T C A C A G T C T A G G T C A C T G A C G T A T A G C G A T C C G T A A C T G T C G A T G C A | 1e-36 | -8.296e+01 | 0.63% | 0.02% | 47.9bp (0.0bp) | IRF4(IRF)/GM12878-IRF4-ChIP-Seq(GSE32465)/Homer(0.733) More Information \| Similar Motifs Found | motif file (matrix) |
| 7 | G T C A G T C A T G A C A C T G A G C T A C T G A C T G A C G T C G A T A C G T | 1e-35 | -8.125e+01 | 0.96% | 0.09% | 51.0bp (42.9bp) | RUNX1(Runt)/Jurkat-RUNX1-ChIP-Seq(GSE29180)/Homer(0.698) More Information \| Similar Motifs Found | motif file (matrix) |
| 8 | C G T A C T A G A T G C C A T G A T C G C G T A C G A T T A G C A C G T G C T A | 1e-34 | -7.948e+01 | 0.61% | 0.02% | 48.8bp (0.0bp) | TFAP4/MA0691.1/Jaspar(0.602) More Information \| Similar Motifs Found | motif file (matrix) |
| 9 | T A C G C T G A C T A G C T G A C T G A A T C G C T G A C T G A C T G A A C G T A C T G C G T A | 1e-33 | -7.603e+01 | 0.59% | 0.03% | 57.1bp (10.5bp) | MAFG::NFE2L1/MA0089.1/Jaspar(0.645) More Information \| Similar Motifs Found | motif file (matrix) |
| 10 | A G C T A C T G A T G C G T A C G C A T A G C T G C A T A C G T G C T A A T G C G A T C A G C T | 1e-28 | -6.456e+01 | 0.65% | 0.06% | 52.2bp (32.4bp) | MYNN(Zf)/HEK293-MYNN.eGFP-ChIP-Seq(Encode)/Homer(0.686) More Information \| Similar Motifs Found | motif file (matrix) |
| 11 | C T A G T C G A C T G A G C T A C T G A T C G A C T G A C G A T C G T A C T A G T G C A C A G T | 1e-28 | -6.456e+01 | 0.65% | 0.05% | 51.3bp (0.0bp) | PB0192.1\_Tcfap2e\_2/Jaspar(0.785) More Information \| Similar Motifs Found | motif file (matrix) |
| 12 | C T G A C G T A C G T A A C G T C G T A C G T A C G T A A C T G A C G T A G C T | 1e-23 | -5.458e+01 | 0.67% | 0.07% | 55.9bp (11.8bp) | PH0075.1\_Hoxd10/Jaspar(0.660) More Information \| Similar Motifs Found | motif file (matrix) |
| 13 | C A G T A C T G C A G T C T A G T C G A A C G T A T G C C G A T C A T G C G A T T A C G A T G C | 1e-23 | -5.346e+01 | 0.57% | 0.05% | 46.8bp (45.3bp) | NR4A2/MA0160.1/Jaspar(0.611) More Information \| Similar Motifs Found | motif file (matrix) |
| 14 | G T C A T C G A C G T A A C T G A C T G T C A G C G T A G T A C C G T A A C T G | 1e-22 | -5.118e+01 | 1.98% | 0.63% | 54.6bp (62.7bp) | PRDM1(Zf)/Hela-PRDM1-ChIP-Seq(GSE31477)/Homer(0.684) More Information \| Similar Motifs Found | motif file (matrix) |
| 15 | G A C T A C G T G T A C C A T G C T A G T C G A A G T C C A T G G T A C C G A T | 1e-21 | -4.970e+01 | 0.43% | 0.03% | 54.1bp (8.0bp) | HINFP/MA0131.2/Jaspar(0.640) More Information \| Similar Motifs Found | motif file (matrix) |
| 16 | C G T A G C T A A G C T C A T G C T G A A T C G C G T A G T A C A G C T C G T A | 1e-21 | -4.905e+01 | 1.57% | 0.44% | 56.5bp (58.6bp) | Pax7(Paired,Homeobox),long/Myoblast-Pax7-ChIP-Seq(GSE25064)/Homer(0.676) More Information \| Similar Motifs Found | motif file (matrix) |
| 17 | T G C A C G T A C A T G A G T C T C A G T G A C T G A C C T A G C T G A T A G C | 1e-20 | -4.811e+01 | 0.53% | 0.05% | 53.2bp (27.4bp) | POL010.1\_DCE\_S\_III/Jaspar(0.597) More Information \| Similar Motifs Found | motif file (matrix) |
| 18 | C G T A T C A G T C G A C T G A C T A G C A T G T G A C G C A T A C G T A G T C G C A T C A T G | 1e-20 | -4.811e+01 | 0.53% | 0.05% | 45.2bp (30.6bp) | POL008.1\_DCE\_S\_I/Jaspar(0.572) More Information \| Similar Motifs Found | motif file (matrix) |
| 19 | A T C G T G A C G T C A C A G T C A T G C A T G C T A G T A C G C G T A C T A G T C A G C T G A | 1e-19 | -4.548e+01 | 0.51% | 0.04% | 40.7bp (33.3bp) | Znf263(Zf)/K562-Znf263-ChIP-Seq(GSE31477)/Homer(0.708) More Information \| Similar Motifs Found | motif file (matrix) |
| 20 | A G C T A G T C A C T G A G T C C G A T A T C G G T A C A T G C C T A G A C G T | 1e-18 | -4.290e+01 | 0.49% | 0.05% | 40.3bp (56.1bp) | POL010.1\_DCE\_S\_III/Jaspar(0.596) More Information \| Similar Motifs Found | motif file (matrix) |
| 21 | A C T G A C T G C G T A C G T A G T C A A C G T A C T G A G T C C G T A A C T G | 1e-18 | -4.290e+01 | 0.49% | 0.05% | 49.9bp (13.0bp) | GFY(?)/Promoter/Homer(0.698) More Information \| Similar Motifs Found | motif file (matrix) |
| 22 | G T A C G C T A T G C A C T G A C T A G G C T A C T G A C G A T A T C G C G T A C G A T G C T A | 1e-18 | -4.290e+01 | 0.49% | 0.05% | 38.4bp (56.3bp) | SOX10/MA0442.2/Jaspar(0.613) More Information \| Similar Motifs Found | motif file (matrix) |
| 23 | C G T A G T C A C G T A A C G T A C T G C G T A A G T C C G T A A C G T A G C T | 1e-17 | -4.052e+01 | 0.37% | 0.03% | 56.4bp (0.0bp) | PB0169.1\_Sox15\_2/Jaspar(0.769) More Information \| Similar Motifs Found | motif file (matrix) |
| 24 | C A T G C A T G C A G T A C T G C A G T A T G C G T A C C A T G T C G A C A G T | 1e-17 | -4.036e+01 | 0.55% | 0.06% | 53.2bp (46.3bp) | PH0164.1\_Six4/Jaspar(0.737) More Information \| Similar Motifs Found | motif file (matrix) |
| 25 | A C T G A G T C A G T C A G C T G T A C A T G C A C G T C T G A A C G T A C T G | 1e-17 | -4.036e+01 | 0.47% | 0.05% | 59.6bp (26.1bp) | Zac1(Zf)/Neuro2A-Plagl1-ChIP-Seq(GSE75942)/Homer(0.694) More Information \| Similar Motifs Found | motif file (matrix) |
| 26 | G A C T A G T C C A G T A T G C T A C G A G C T C G T A A T C G C G A T T A G C | 1e-16 | -3.810e+01 | 0.53% | 0.07% | 52.0bp (38.8bp) | PB0106.1\_Arid5a\_2/Jaspar(0.620) More Information \| Similar Motifs Found | motif file (matrix) |
| 27 | A T G C T G A C G T A C C G T A T A C G T C G A T A G C T C A G A C G T A T G C G C A T C T A G | 1e-16 | -3.785e+01 | 0.45% | 0.06% | 48.2bp (51.4bp) | Smad4(MAD)/ESC-SMAD4-ChIP-Seq(GSE29422)/Homer(0.695) More Information \| Similar Motifs Found | motif file (matrix) |
| 28 | A G T C A C T G T G A C A C T G C G A T A G C T A G T C A G T C G T A C C T A G | 1e-16 | -3.756e+01 | 0.35% | 0.01% | 46.1bp (0.0bp) | TFDP1/MA1122.1/Jaspar(0.722) More Information \| Similar Motifs Found | motif file (matrix) |
| 29 | A G T C A C T G C T G A C G A T C T A G C G T A A C G T C A T G G T C A A C G T | 1e-16 | -3.756e+01 | 0.35% | 0.03% | 39.3bp (38.6bp) | PH0017.1\_Cux1\_2/Jaspar(0.700) More Information \| Similar Motifs Found | motif file (matrix) |
| 30 | A T C G A C T G C G T A G C T A C G T A G T C A C G T A A G T C | 1e-15 | -3.627e+01 | 3.82% | 2.01% | 53.5bp (56.7bp) | NFATC2/MA0152.1/Jaspar(0.792) More Information \| Similar Motifs Found | motif file (matrix) |
| 31 | T A C G C T G A T C A G T G C A T G C A G C T A C A T G A G C T T G A C G T A C | 1e-14 | -3.339e+01 | 0.61% | 0.10% | 53.7bp (42.6bp) | PRDM1/MA0508.2/Jaspar(0.730) More Information \| Similar Motifs Found | motif file (matrix) |
| 32 | A C G T A C T G C G T A C G T A A C T G A C G T A G T C A C T G | 1e-13 | -3.180e+01 | 0.31% | 0.03% | 40.8bp (32.7bp) | MF0002.1\_bZIP\_CREB/G-box-like\_subclass/Jaspar(0.674) More Information \| Similar Motifs Found | motif file (matrix) |
| 33 | A C G T C G T A A C G T A G T C A C G T A C G T A C G T C G A T A C G T A C T G | 1e-13 | -3.157e+01 | 0.47% | 0.07% | 55.3bp (68.6bp) | SD0003.1\_at\_AC\_acceptor/Jaspar(0.649) More Information \| Similar Motifs Found | motif file (matrix) |
| 34 | A G T C C A T G C G A T A C G T A G T C A C T G A C T G A T G C | 1e-13 | -3.060e+01 | 0.39% | 0.04% | 47.5bp (10.9bp) | ZBTB7A/MA0750.2/Jaspar(0.589) More Information \| Similar Motifs Found | motif file (matrix) |
| 35 | C T A G A G T C G T A C C G T A C G T A A G T C A C T G A C T G | 1e-13 | -3.038e+01 | 0.63% | 0.13% | 47.5bp (53.1bp) | AMYB(HTH)/Testes-AMYB-ChIP-Seq(GSE44588)/Homer(0.739) More Information \| Similar Motifs Found | motif file (matrix) |
| 36 | C G T A G T A C G C T A C T A G C A G T T G A C T A C G A C G T | 1e-12 | -2.961e+01 | 9.06% | 6.42% | 56.0bp (52.6bp) | MYF6/MA0667.1/Jaspar(0.722) More Information \| Similar Motifs Found | motif file (matrix) |
| 37 | C G T A A C G T C G T A A G C T C G T A G T A C A C T G A C G T T C G A A C G T | 1e-12 | -2.828e+01 | 0.37% | 0.05% | 45.9bp (0.0bp) | CLOCK/MA0819.1/Jaspar(0.684) More Information \| Similar Motifs Found | motif file (matrix) |
| 38 \* | A C G T A C T G A G T C A C T G A C G T C T A G A G T C A C G T | 1e-11 | -2.683e+01 | 0.88% | 0.26% | 54.0bp (37.2bp) | Arnt:Ahr(bHLH)/MCF7-Arnt-ChIP-Seq(Lo\_et\_al.)/Homer(0.849) More Information \| Similar Motifs Found | motif file (matrix) |
| 39 \* | A T C G A C G T A G T C C G T A A G T C A C T G A C G T A C T G | 1e-11 | -2.551e+01 | 0.82% | 0.23% | 46.1bp (79.8bp) | Arntl/MA0603.1/Jaspar(0.977) More Information \| Similar Motifs Found | motif file (matrix) |
| 40 \* | C G T A C G T A A T C G C T A G A G T C A G T C A C T G A C G T | 1e-10 | -2.449e+01 | 0.51% | 0.11% | 35.9bp (31.7bp) | MYB(HTH)/ERMYB-Myb-ChIPSeq(GSE22095)/Homer(0.670) More Information \| Similar Motifs Found | motif file (matrix) |
| 41 \* | G A C T C G T A A C T G T G C A A C G T C G A T C T A G G T C A A C G T C G A T A C T G G T C A | 1e-10 | -2.363e+01 | 0.25% | 0.02% | 53.5bp (2.2bp) | PBX1/MA0070.1/Jaspar(0.682) More Information \| Similar Motifs Found | motif file (matrix) |
| 42 \* | A G T C A G T C A C T G A C T G A G T C A C G T A C G T A C T G | 1e-10 | -2.363e+01 | 0.25% | 0.03% | 72.7bp (47.8bp) | TFCP2/MA0145.3/Jaspar(0.654) More Information \| Similar Motifs Found | motif file (matrix) |
| 43 \* | A T G C C G T A A T G C C G T A T A G C C T G A A G T C C T G A | 1e-9 | -2.282e+01 | 12.85% | 10.09% | 49.7bp (62.6bp) | PB0130.1\_Gm397\_2/Jaspar(0.755) More Information \| Similar Motifs Found | motif file (matrix) |
| 44 \* | C G T A C G T A A C T G C G T A C G T A A C T G C G T A A C T G A C T G C G T A | 1e-9 | -2.163e+01 | 0.31% | 0.05% | 42.4bp (7.6bp) | SPI1/MA0080.4/Jaspar(0.658) More Information \| Similar Motifs Found | motif file (matrix) |
| 45 \* | A C T G A C G T A C G T C G T A C G T A C G T A A C G T A G T C | 1e-8 | -2.027e+01 | 0.59% | 0.17% | 63.6bp (49.7bp) | Gfi1/MA0038.1/Jaspar(0.707) More Information \| Similar Motifs Found | motif file (matrix) |
| 46 \* | A T C G A C T G A C T G A C G T A C T G A C T G C T A G A G T C A T C G A C T G | 1e-8 | -1.953e+01 | 0.29% | 0.04% | 59.4bp (27.6bp) | Egr2(Zf)/Thymocytes-Egr2-ChIP-Seq(GSE34254)/Homer(0.841) More Information \| Similar Motifs Found | motif file (matrix) |
| 47 \* | A C G T C A G T C G A T A G T C A T G C A C T G A C T G C G T A | 1e-8 | -1.900e+01 | 0.53% | 0.15% | 51.5bp (54.7bp) | Elk4(ETS)/Hela-Elk4-ChIP-Seq(GSE31477)/Homer(0.796) More Information \| Similar Motifs Found | motif file (matrix) |
| 48 \* | G C A T G T A C A C G T C T G A A G C T G T C A G C A T G T C A C A T G T G C A C G A T G C T A | 1e-8 | -1.893e+01 | 1.60% | 0.79% | 38.7bp (62.9bp) | PB0198.1\_Zfp128\_2/Jaspar(0.747) More Information \| Similar Motifs Found | motif file (matrix) |
| 49 \* | C G T A C G T A A C G T C G T A A C G T A C G T A C T G A C T G | 1e-7 | -1.766e+01 | 3.31% | 2.11% | 56.2bp (58.9bp) | Arid5a/MA0602.1/Jaspar(0.878) More Information \| Similar Motifs Found | motif file (matrix) |
| 50 \* | A C G T A G T C A C T G C G T A A C G T A G T C A G T C A C G T | 1e-7 | -1.749e+01 | 0.27% | 0.05% | 43.4bp (40.9bp) | HNF6(Homeobox)/Liver-Hnf6-ChIP-Seq(ERP000394)/Homer(0.732) More Information \| Similar Motifs Found | motif file (matrix) |
