## Supplementary Materials for "Germline and somatic genetic effects on gene expression and outcome in patients with Multiple Myeloma"

### Assessing the quality of germline variant and genotype calls

Concordance of chr22 variant and genotype calls between low-coverage Whole Genome Sequence and high-coverage Whole Exome Sequence data was assessed with VCFtools (Danecek et al. 2011) vcf-compare. While the alternative alleles were called with a lower rate from the low-coverage WGS data, the overall concordance was high, showing that germline genotypes can be reliably called from low-coverage WGS data (Supplementary Table 1, 2). Sample-wise genotype concordance varied from 87.13% to 95.78% with a mean of 93.22%.

**Table 1.** Variant call concordance between WGS and WES data

| | Reference allele | Alternative allele |
|---|---|---|
| Matches | 44,858 | 43,812 |
| Mismatches | 412 | 1046 |
| Total calls | 45,270 | 44,858 |
| Concordance | 99.09% | 97.67% |

**Table 2.** Genotype call concordance between WGS and WES data.

| | REF/REF | REF/ALT | ALT/ALT |
|---|---|---|---|
| Mismatches | 352,360 | 100,598 | 82,056 |
| Matches | 8,408,336 | 1,175,975 | 858,908 |
| Concordance | 95.98% | 92.12% | 91.28% |

### Assessing the quality of somatic variant calls

Variant concordance was calculated between the high-coverage whole-exome sequenced and low coverage whole-genome sequenced data for 513 of the 520 samples in the somatic data. 7 of the 520 samples did not undergo whole-exome sequencing, so variant concordance was not calculated for these samples. The workflow for calculating variant concordance was as follows:

first, the number of mutations called in both the somatic whole-genome and somatic whole-exome sequenced data was obtained; second, the number of mutations that were called in the somatic whole genome data but not the somatic whole exome data was obtained; third, the coverage data was compared for the mutations that were called in the somatic whole-genome sequenced but not the somatic whole-exome sequenced data using bam-readcounts (McDonnell Genome Institute. 2018); fourth, the number of mutations that were called in whole-genome that had concordant coverage with the whole-exome sequenced data, with whole-exome sequenced data coverage greater than 5 was obtained. The formula to calculate variant concordance was as follows:

$$\text{Variant concordance} = \frac{(N \text{ of calls in both whole exome and whole genome data}) + (N \text{ of concordant calls from coverage analysis})}{(N \text{ of calls in both whole exome and whole genome data}) + (N \text{ of calls in whole genome, not the whole exome, with whole exome coverage greater than or equal to } 5x)}$$

Across the 513 patients, the average variant concordance was 75.15% with a standard deviation of 12.55%, suggesting that somatic genotypes can be called from low coverage whole-genome sequenced data.

### Evaluation of genotype imputation accuracy

Imputation accuracy was evaluated by the internal cross-validation performed by IMPUTE2 (Howie, Donnelly, and Marchini 2009). After filtering for genotype probability score ≥0.9, concordance between measured and imputed genotypes varied from 89.3 to 99.1%.

### Accounting for technical batch effects in gene expression data

Accounting for technical and other confounding sources of expression variation may greatly increase statistical power in association analyses. We used a probabilistic estimation of expression residuals (Stegle et al. 2012) to infer hidden sources of variation in expression data. These latent factors were used as surrogate variables for unknown technical batch effects and included as covariates in downstream analyses. 100 hidden factors were included in the model, as recommended in Stegle et al. 2012 (for technical details, see Stegle et al. 2010).

### Accounting for population structure

Population structure may lead to an inflation of *p*-values in association studies (Price et al. 2006). To account for population structure in the eQTL analyses, we used the R package *SNPRelate* (Zheng et al. 2012) to perform principal component analysis (PCA) on the germline genotype data and included 5 PCs as covariates. To evaluate whether we had successfully accounted for population structure in the eQTL analysis, we used a shuffled expression matrix to create a null-distribution of *p*-values with and without genotype-based PCs (Fig. 1A).

### Evaluating the effect of Copy Number Alterations in eQTL calling

Copy number changes may alter gene expression levels. To evaluate the impact of tumor copy number alterations (CNAs) on eQTL calls, we first obtained CNV information from 569 samples collapsed to gene levels. Then, using chr22, we fitted a linear model between the CNV and gene expression levels and used the residuals of this model to repeat the eQTL mapping using this subset of 569 samples.

### Identifying puromycin concentration to select for successful transformants

To identify the optimal puromycin concentration to select for the transformed cells containing gRNA CROP-seq vector, we set up a killing curve experiment. Briefly, cells were collected by centrifuging at 300g for 5 minutes, then dissolved in AR5 media without any antibiotics to a concentration of 400,000 cells per ml. The experiment was performed on a 96 well plate with 1ml of cell culture per well, and various puromycin concentration was added into the cell culture (Supplementary Figure 1). Cells were then incubated in the incubator at $37^0$C for three days, then the cells were stained with CellTiter Glo (Promega) and fluorescence intensity was measured using the Cytation 3 imaging reader (BioTek). Two replicates were performed per cell line and the optimal puromycin was determined as the lowest concentration that killed all the non-transformed cells.

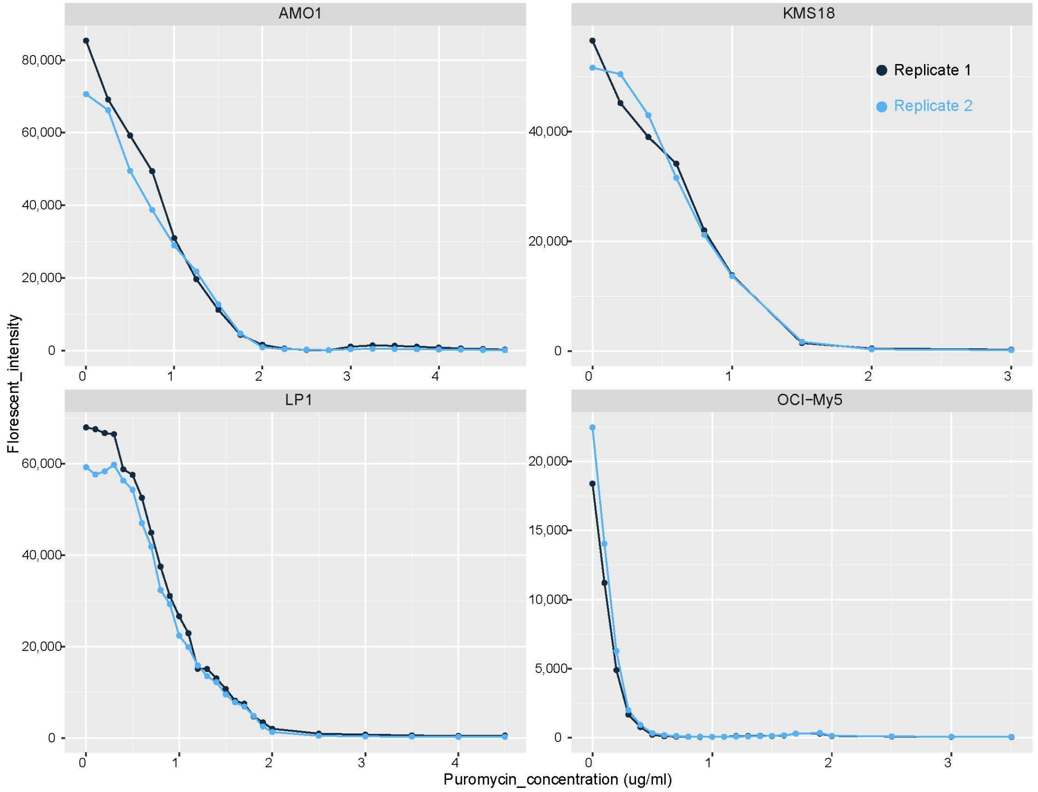


**Supplementary Figure 1**. Killing curve experiment for the cell lines used in the study
